# Reactive Inhibitory Control Precedes Overt Stuttering Events

**DOI:** 10.1101/2022.08.02.501857

**Authors:** Joan Orpella, Graham Flick, M. Florencia Assaneo, Liina Pylkkänen, David Poeppel, Eric S. Jackson

**Author notes:** Corresponding authors Joan Orpella, Eric S. Jackson.

## Abstract

Research points to neurofunctional differences underlying fluent speech between stutterers and non-stutterers. Considerably less work has focused on processes that underlie stuttered vs. fluent speech. Additionally, most of this research has focused on speech motor processes despite contributions from cognitive processes that occur prior to the onset of stuttered speech. We used MEG to test the hypothesis that reactive inhibitory control is triggered prior to stuttered speech. Twenty-nine stutterers completed a delayed-response task that featured a cue (prior to a go cue) signaling the imminent requirement to produce a word that was either stuttered or fluent. Consistent with our hypothesis, we observed increased beta power in the R-preSMA –an area implicated in reactive inhibitory control– in response to the cue preceding stuttered vs. fluent productions. Beta power differences between stuttered and fluent trials correlated with stuttering severity and participants’ percentage of trials stuttered increased exponentially with beta power in the R-preSMA. Trial-by-trial beta power modulations in the R-preSMA following the cue predicted whether a trial would be stuttered or fluent. Stuttered trials were also associated with delayed speech onset suggesting an overall slowing or freezing of the speech motor system that may be a consequence of inhibitory control. Post-hoc analyses revealed that independently-generated anticipated words were associated with greater beta power and more stuttering than researcher-assisted anticipated words, pointing to a relationship between self-perceived likelihood of stuttering (i.e., anticipation) and inhibitory control. This work offers a neurocognitive account of stuttering by characterizing the cognitive processes that precede overt stuttering events.

## Introduction

Stuttering is typically characterized by its symptoms including overt interruptions in speech (i.e., part-word repetitions, blocks, and prolongations). However, research suggests that stuttering events begin prior to speech onset, which is consistent with the experience of stutterers [1–4]. Several studies have used imaging techniques with high temporal resolution, including MEG and EEG, revealing distinct neural activity preceding speech onset in adults who stutter compared to adults who do not stutter [5–9]. Vanhoutte et al. [9], for instance, observed heightened contingent negative variation (CNV), a slow-rising scalp potential associated with speech initiation and basal ganglia-thalamocortical (BGTC) loop activity prior to speech onset. In addition, stutterers exhibit increased suppression of beta oscillations in mouth motor areas prior to speech onset, taken to indicate a lack of coordination in the BGTC loop that hampers speech initiation and sequencing [5,6]. Stutterers also exhibit reduced auditory evoked potentials compared to non-stutterers prior to speech onset, suggesting a reduced capacity to modulate auditory cortical activity during speech preparation [10,11]. This points to inefficient forward modeling which could potentially lead to overt stuttering [12,13]. Together, these studies reveal divergent neural activity prior to speech onset in stutterers compared to typically fluent speakers.

Fewer studies have explored brain activity preceding overtly stuttered vs. fluent speech, largely due to the difficulty of eliciting stuttering reliably during neuroimaging. For instance, Sowman et al. [14] investigated stuttered versus fluent productions of a single vowel in one participant and found reduced activation in regions associated with articulatory planning suggesting that stutterers experience planning difficulties related to stuttered but not fluent speech. Similarly, Vanhoutte et al. [15] reported a reduced CNV preceding stuttered vs. fluently produced words in seven participants. They argued that a reduced CNV in left hemisphere speech regions reflects atypical speech initiation processes that result in overt stuttering [15], whereas their previous finding of elevated CNV prior to fluent speech in stutterers reflects compensatory mechanisms particularly in right hemisphere regions [9]. Mersov et al. [16] did not observe significant differences preceding stuttered versus fluent productions using MEG but found a weak trend for delayed motor beta suppression preceding stuttered speech. In all, the available data suggest that neural activity preceding stuttered speech differs from that preceding fluent speech.

While the above studies focused on speech motor processes, recent work has begun to explore cognitive processes during speech preparation that influence how stuttering manifests itself overtly [17]. For example, stutterers develop the ability to anticipate stuttering or predict that a particular word will be stuttered if the word is produced as planned [18,19]. Stuttering anticipation develops through associative learning whereby stuttered words or sounds become linked to negative reactions from listeners or the environment such that stutterers become averse to overt stuttering events [20,21]. Critically, adult stutterers anticipate stuttering with high accuracy [> 85%; 20–22] suggesting that it is highly likely that a stuttered word would have been anticipated given that there is enough time between knowing a word will need to be produced and producing it, and also given that attention is directed towards speech [18]. This is particularly the case in experimental contexts during which participants produce single words [17].

It is likely that an inhibitory response is triggered prior to overtly stuttered speech due to the negative consequences that stutterers associate with overt stuttering [18,20,21] and their ability to predict it [22–24]. Neural evidence for such a response comes from a recent investigation by Jackson et al. [17] which revealed increased right dorsolateral prefrontal cortex (R-DLPFC) activity seconds prior to producing anticipated vs. unanticipated words [17]. This activity is consistent with a *proactive* inhibitory control response to stuttering anticipation. Proactive inhibitory control is a forward-looking process aimed at preventing or delaying undesired actions and associated with heightened R-DLPFC activity [25,26]. Overt stuttering is an undesired action because of its associated negative consequences. Therefore, individuals who stutter employ proactive inhibitory control to cope with impending stuttering, and will often implement learned strategies such as modifying the speech plan to circumvent the anticipated word (e.g., by stalling or switching a word) [18,27,28]. This is consistent with the finding of delayed speech initiation times for anticipated words in Jackson et al. (2022).

It has also been proposed that *reactive* inhibitory control plays a role in stuttering [29–31]. In contrast to proactive inhibitory control, reactive inhibitory control refers to an automatic and rapid response triggered by an external cue to halt a planned action. Reactive inhibitory control is typically tested using stop signal tasks during which participants’ neural responses to external signals to stop an initiated action are measured. The most consistent neural marker of reactive inhibition is increased beta power in regions of the action-stopping network, including the right pre-supplementary motor area (R-preSMA), right inferior frontal gyrus (R-IFG), and subthalamic nucleus [26,32]. Unlike proactive inhibitory control which may arise when a stutterer thinks about having to produce an anticipated word, reactive inhibitory control may be more immediate and triggered by an external cue that signals that overt stuttering is imminent. Consider a situation in which a stutterer is going to introduce themself to a new person. As they approach the new person, the stutterer knows they will have to say their name, which is an anticipated word for many stutterers. Leading up to production of the word, the stutterer may delay speech initiation while thinking of strategies to circumvent overt stuttering such as using a starter utterance, e.g., “my name is” or, “um.” These behaviors are likely reflective of proactive inhibitory control given their prospective nature. At the same time, the stutterer may exhibit reactive inhibitory control, for example, when the new person extends their hand and introduces themself because, at this moment, the stutterer knows that they will have to produce their name imminently. In this example, the gesture of the new person (e.g., the extended hand) serves as an external cue that the stutterer will soon have to produce their name, an action to which the stutterer is averse. This cue may trigger a “freezing” response to the threat of producing the anticipated word [33]. Due to its immediacy and automaticity, the cue indicating that the speaker will have to produce their name imminently likely triggers reactive rather than proactive inhibitory control.

In the current study, we investigated reactive inhibitory control responses prior to overt stuttering events using MEG. MEG is an ideal functional neuroimaging technique to study modulations in beta oscillatory activity which are a hallmark characteristic of reactive inhibitory control [26,34,35]. We employed our recently introduced method to elicit a semi-balanced amount of stuttered and fluent speech across participants [36,37]. Fifty participant-specific anticipated words (i.e., words identified as likely to be stuttered) were determined during a clinical interview procedure in Visit 1. In Visit 2, participants completed a delayed-response task in which they produced these words while neural activity was recorded with MEG. Critically, the go signal in this task was preceded by a cue that informed participants of the imminent requirement to produce the word. We hypothesized that reactive inhibitory control would be triggered by this cue and would be greater prior to overtly stuttered vs. fluent productions. That is, the cue that signals the imminent requirement to speak will act as an implicit no-go signal because of the stutterer’s learned association between stuttering and its negative consequences. A reactive inhibitory control response to the cue would be reflected by increased beta oscillatory activity for stuttered vs fluent trials in the R-preSMA and/or R-IFG. Each participant’s word list consisted of both *independently-generated* anticipated words (i.e., words independently identified by participants as likely to be stuttered) and *researcher-assisted* anticipated words (i.e., words identified by participants as likely to be stuttered with researcher assistance). This allowed for a post-hoc analysis to explore the relationship between anticipation and reactive inhibitory control based on the assumption that independently-generated words are associated with greater anticipation than researcher-assisted words.

## Materials and methods

This study was approved by the Institutional Review Board at New York University. Written consent was obtained from all participants in accordance with the Declaration of Helsinki.

### Participants

Participants were recruited via the last author’s database of stutterers, a mass email from the National Stuttering Association, and word of mouth. Participants included 31 adults who stutter (8 female), with a mean age of 30.1 (SD = 7.8), who all reported a negative history of concomitant neurological, psychiatric, or psychological disorders. All participants reported being native English speakers. Stuttering diagnosis was made by the last author, a speech-language pathologist (SLP) with more than 15 years of experience and expertise in stuttering intervention. All participants also self-reported as a stutterer and exhibited three or more stuttering-like disfluencies [38] with temporally aligned physical concomitants (e.g., eye blinking, head movements) during a 5-10 minute conversation. The Stuttering Severity Index - 4th Edition (SSI-4) [39] was also administered, and participants responded to three subjective severity questions (Q1-Q3) via 5-pt. Likert scales: Q1) How severe would you rate your stuttering?; Q2) How severe would other people rate your stuttering?; Q3) Overall, how much does stuttering impact your life? (1 = Mild, 5 = Severe). There were two visits. Visit 1 included diagnostic testing and a clinical interview to determine participant-specific stimuli, i.e., anticipated words. Visit 2 included MEG testing.

### Clinical Interview

The interview was adapted from Jackson et al. [37]. In that study, both anticipated and unanticipated words were elicited from participants, which yielded a near equal distribution of stuttered and fluent speech during fNIRS recording [17]. However, Jackson et al. [17,37] included interactive speech whereas the current study does not; the likelihood of stuttering during testing with face-to-face communication is higher. Therefore, we only elicited anticipated words in the current study to increase the probability of a near-balanced distribution of stuttered and fluent speech in the absence of face-to-face communication (i.e., in the shielded room while MEG data were recorded). The interview is fully described in Jackson et al. [37], but is also summarized here. Participants were initially asked if they anticipated stuttering; all participants confirmed that they did. Participants were then asked to identify words that they anticipate stuttering on. Most participants identified at least a few words, though there was variability across participants (as in Jackson et al.) Participants were also provided with prompts (e.g., “What about your name?”, “Is your job title a difficult word for you?”). Participants were then asked about anticipated sounds, i.e., word-initial sounds that are problematic, which were used to create additional anticipated words beginning with these sounds. This was done ultimately to produce a list of 50 different, participant-specific words to be presented during MEG testing (visit 2). Critically, all 50 words were anticipated, regardless of how they were elicited.

### Stimuli

Each participant had their own list of 50 anticipated words, which were presented during the MEG experiment. While all of these words were considered anticipated words, there were words that were identified by the participant independently (i.e., *independently-generated* words), and words that were generated with researcher assistance (i.e., *researcher-assisted* words). Researcher-assisted words could be identified, for example, by asking participants whether specific words are difficult or asking participants about difficult sounds, and then identifying words that began with these difficult sounds. Researcher-assisted words were developed using an online word generator and were five syllables in length, because longer words tend to be stuttered more than shorter words [40]. For example, if /b/ was identified as an anticipated sound, words like *biochemistry*, *biological,* and *biographical* may have been included. Independently-generated words were typically shorter than researcher-assisted words. Given that anticipation is tightly linked to an individual’s history with stuttering [20,21], we assumed that independently-generated words elicit greater stuttering anticipation than researcher-assisted words. Independently-generated and researcher-assisted anticipated words were each produced six times during the task.

### Task

The behavioral task is depicted in Fig. 1. Each trial began with a fixation cross (baseline period) of variable duration (1 – 1.5 sec). A word from the anticipated words list (see Stimuli section) appeared in the center of the screen (0.4 sec) followed by a blank screen of variable duration (0.4 – 0.8 sec). After this blank screen, there was either a *speak* trial or a *catch* trial. For speak trials, a white asterisk appeared (henceforth, cue), signaling the requirement to speak the word at the onset of the following green asterisk (henceforth, go-signal). The duration of the white asterisk was 0.2 seconds and was always followed by 0.8 seconds of blank screen. The time between the onsets of the white asterisk (cue) and the green asterisk (go signal) was therefore always 1 second. The duration of the green asterisk on the screen was 0.5 seconds and was followed by a variable blank period (2 – 3 sec) to allow the participant to produce the word. The word STOP appeared at the end of this blank period, requiring participants to abort any incomplete speech acts and prepare for the next trial by remaining as still as possible. Catch trials (15% of trials) were introduced to create uncertainty about the requirement to speak the anticipated word. In these catch trials, a red asterisk followed the blank screen and the participant was required to remain silent and await the following trial (Fig. 1). The overall design thus simulates a common experience of people who stutter–the stutterer is required to produce an anticipated word (e.g., one’s own name) within an expected time window (e.g., after the conversational partner extends their hand and says “What is your name?”). The critical window for analysis was the cue period of 1 sec before the go signal (i.e., between the white and the green asterisks). The task consisted of six blocks; each block included the list of 50 words presented in randomized order, for a total of 300 words per MEG session. Participants’ faces and speech during the experiment were recorded.

**Figure 1.**
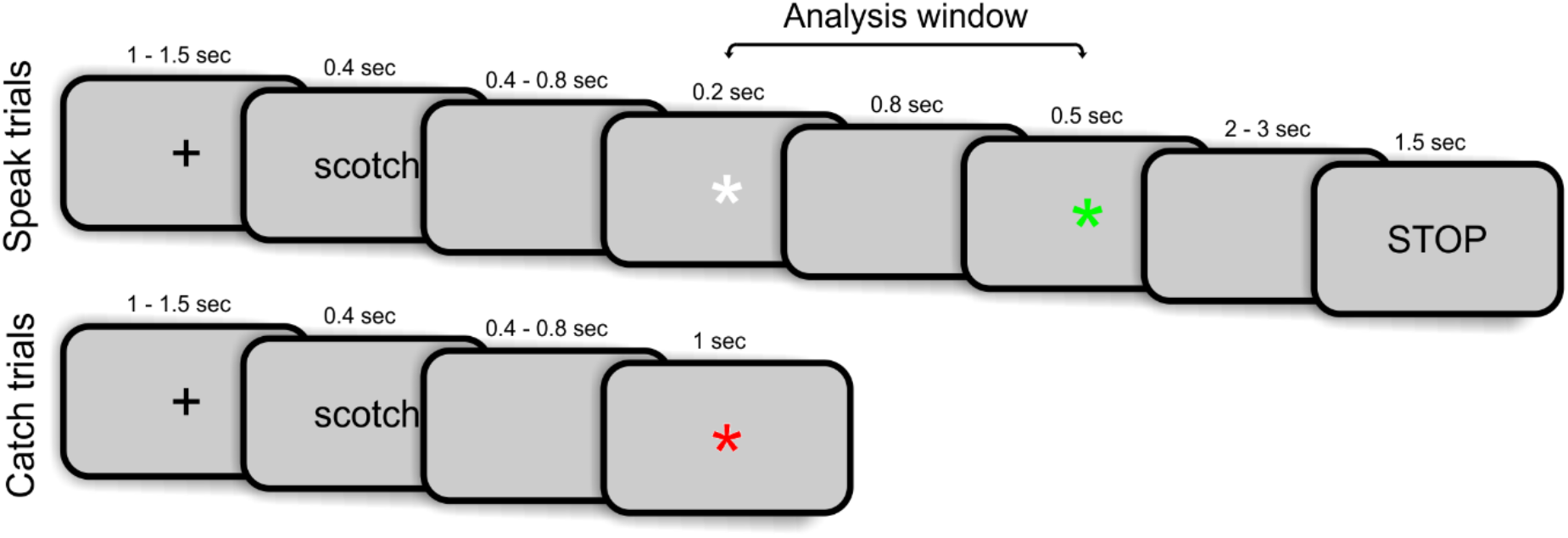
Behavioral task. Each trial began with a fixation cross of variable duration (Baseline period). Stimulus words appeared in the center of the screen followed by a blank screen of variable duration. For speak trials, a white asterisk appeared (cue), signaling the requirement to speak the word on the following green asterisk (go signal). Participants had 2 – 3 s to produce the words. Catch trials started in the same manner, however, a red asterisk appeared after the initial blank screen, indicating that participants should remain silent until the next trial.

### MEG data acquisition and preprocessing

Neuromagnetic responses were acquired using a 157-channel whole-head axial gradiometer (Kanazawa Institute of Technology, Japan) situated in a magnetically shielded room, with a sampling rate of 1000Hz. To monitor head position during the recordings, five electromagnetic coils were attached to the subject’s head. We registered the location of these coils with respect to the MEG sensors before and after each block of the experiment. Participants’ head shape was digitized immediately before the MEG recordings using a Polhemus digitizer and 3D digitizer software (Source Signal Imaging) along with 5 fiducial points, to align the position of the coils with participants’ head shape, and 3 anatomical landmarks (nasion, and bilateral tragus), to further allow for the co-registration of participant’s MEG data with an anatomical MRI template. An online band-pass filter (1Hz-200Hz) was applied to all MEG recordings.

Data preprocessing was conducted using custom Python scripts and MNE-python software [41]. Bad channels were first selected and interpolated using spherical spline interpolation. A least-squares projection was then fitted to the data from a 2-minute empty room recording acquired at the beginning of each MEG session and the corresponding component was removed. MEG signals were next digitally low-pass filtered at 50Hz using MNE-python’s default parameters with firwin design and finally epoched between −1700 ms and 1500 ms relative to the onset of presentation of the cue (white asterisk; Fig. 1). Linear detrending was applied to the epochs to account for signal drift. Baseline correction was applied at the analysis phase (see below). An independent component analysis was used to correct for cardiac, ocular, and muscle artifacts. The epochs resulting from these steps were visually inspected and remaining artifactual trials were discarded from further analysis.

### Data Analysis - Behavioral

Trials were judged to be stuttered, fluent, or errors by the last author, an SLP with more than 15 years of experience and expertise in stuttering intervention. Thirty percent of the trials (randomly selected) were also classified by another SLP, blind to the study, to determine interrater reliability. Stuttered trials were those with stuttering-like disfluencies including blocks, prolongations, or part-word repetitions. Error trials were those in which participants forgot or did not attempt to produce the target word. Speech onset time was calculated as the time between the go signal (i.e., the green asterisk) and speech onset as defined by the first articulatory movement or accessory behavior [as in 17]). Accessory behaviors refer to non-speech behaviors that often co-occur with stuttering events (e.g., eye blinking, facial tension). Articulatory onset was defined as the first articulatory movement, which was determined based on visual inspection using Davinci Resolve (Black Magic Design, Australia). This procedure allowed for frame-by-frame scanning (29.97 frames per second) of the recordings of participants’ faces. A generalized linear mixed-effects model fit by maximum likelihood (family = binomial) in R[42] was used to assess variables that contributed to stuttered speech. Fixed factors included stimulus type (independently-generated or researcher-assisted), word length (number of letters), initial phoneme (consonant or vowel), and trial number (of experiment). Participant was included as a random effects factor and random slopes for stimulus by participant were also included. Multiple linear regression was used to assess whether SSI-4 score, the number of independently generated anticipated words, and stuttering impact (Q3) contribute to the percentage of trials stuttered for each participant. Finally, a linear mixed-effects model was used to assess potential factors impacting speech onset time. Speech outcome was a fixed factor and participant was a random factor.

### Data Analysis - MEG

#### Time-frequency analysis in sensor space

To determine differences in beta power between stuttered and fluent trials, we conducted a time-frequency analysis. Prior to the decomposition, trial types were equalized in counts using MNE-python’s equalize_epoch_counts function to avoid biases in the time-frequency decomposition toward either condition (*stuttered, fluent*) [41]. In brief, an equal number of trials from each condition is selected while minimizing the differences in the event times of the two conditions to prevent differences due to time-varying characteristics in the data (e.g., adaptation, noise). For the decomposition, we used a Stockwell transform [43] with a Gaussian window of width = 0.5, which offers a good balance between temporal and frequency resolution. The decomposition was performed for each condition separately (*stuttered*, *fluent*), for frequencies between 12 and 30 Hz (beta frequency band) and times between the cue presentation and the presentation of the go signal (Fig. 1). A baseline correction was applied at this stage by subtracting the mean of the values between −1.6 and −1.3 prior to the cue presentation (i.e., within the fixation cross period; Fig. 1) followed by dividing by the mean of the same values (i.e., yielding percent change from baseline). Note that due to the jitter following word presentation, the baseline period differed for each trial. The contrast between *stuttered* and *fluent* trials was performed by subtraction (*stuttered* minus *fluent*) within participant before averaging the resulting difference across participants. To determine the statistical significance of observed differences, we first entered the difference data for each participant into a cluster-based permutation test across participants [44] (one-tailed; 1,000 permutations) using MNE-python’s spatio-temporal_cluster_1samp_test function [41], with an initial *t*-threshold of 2.05 (corresponding to *p* < 0.05 for the number of participants) and a subsequent cluster threshold of *p <* 0.05. The same analysis was performed as a control to determine differences in the theta (4 – 8 Hz), alpha (9 – 12 Hz), and low gamma (30 – 50 Hz) bands during the same time window (cue presentation to go signal).

#### Correlation between overall beta power and behavioral measures

Spearman’s Rank correlation coefficient was used to evaluate the relationship between the beta power difference between stuttered and fluent trials across sensors and stuttering severity. Specifically, we used participants’ beta power difference between stuttered and fluent trials within 0.1 and 0.5 seconds and 22 to 27 Hz and their SSI-4 score (Table 1). One participant had missing data for the SSI-4 and was excluded from this analysis. Statistical significance was established by a threshold of *p* < 0.05. An outlier was removed from this analysis in exceeding 2 SD from the mean of the beta power differences within the identified cluster. We also assessed the correlation between beta power difference and participants’ number of independently-generated words. Statistical significance was established by a threshold of *p* < 0.05 in all cases.

**Table 1.** Behavioral data per participant. Percent (%) trials stuttered includes all trials except catch trials and errors. Independently-generated words include anticipated words identified by participants without any prompting from the researcher (out of 50). These words were coded a posteriori by reviewing the videos for Visit 1. Complete videos were not available for participants 1613 and 1614 and therefore the # of independently words could not be generated for these participants. Visit 1 also included speech sample data for the SSI-4 (Stuttering Severity Instrument – 4^th^ Edition) and impact questions: Q1: How severe would you rate your stuttering?; Q2: How severe would other people rate your stuttering?; Q3: Overall, how much does stuttering impact your life? (1 = Mild, 5 = Severe). Speech sample data were not available for 1613 but were for 1614. Reaction time was calculated using Visit 2 videos. Location of the go signal was not available for 1611 and 1619-1621. ms = milliseconds. Due to researcher error, participants 1607-1621 completed a 7-point Likert scale for Q1-Q3, which was subsequently normalized to a 5-point scale.

| ID | % Trials stuttered | # of independently-generated words | SSI-4 | SSI-4 severity | Q1 | Q2 | Q3 | Reaction time (ms) |
| --- | --- | --- | --- | --- | --- | --- | --- | --- |
| 1607 | 46% | 50 | 22 | Mild | 1.43 | 3.57 | 4.29 | 258.62 |
| 1609 | 63% | 12 | 30 | moderate | 3.57 | 3.57 | 2.86 | 289.65 |
| 1610 | 67% | 14 | 20 | mild | 0.71 | 0.71 | 1.43 | 383.42 |
| 1611 | 78% | 3 | 32 | severe | 4.29 | 3.57 | 2.14 |  |
| 1613 | 22% |  |  |  |  |  |  | 473.85 |
| 1614 | 32% |  | 19 | mild | 2.14 | 2.86 | 3.57 | 240.26 |
| 1615 | 48% | 3 | 31 | moderate | 2.14 | 2.86 | 4.29 | 737.81 |
| 1619 | 10% | 8 | 16 | very mild | 2.14 | 3.57 | 5.00 |  |
| 1620 | 5% | 4 | 26 | moderate | 2.86 | 3.57 | 2.14 |  |
| 1621 | 68% | 1 | 30 | moderate | 2.86 | 2.14 | 1.43 |  |
| 1634 | 14% | 2 | 37 | very severe |  |  |  | 700.10 |
| 1636 | 65% | 9 | 28 | moderate | 5 | 4 | 4 | 344.04 |
| 1637 | 7% | 9 | 19 | mild | 2 | 2 | 3 | 643.71 |
| 1675 | 50% | 0 | 35 | severe | 4 | 4 | 2 | 711.11 |
| 1678 | 3% | 8 | 10 | very mild | 2 | 3 | 2 | 448.16 |
| 1679 | 5% | 9 | 10 | very mild | 1 | 3 | 3 | 634.70 |
| 1681 | 6% | 14 | 13 | very mild | 1 | 1 | 2 | 390.76 |
| 1686 | 48% | 6 | 17 | very mild | 1 | 3 | 5 | 1071.84 |
| 1687 | 36% | 9 | 18 | mild | 3 | 4 | 5 | 833.58 |
| 1690 | 33% | 7 | 25 | moderate | 2 | 1 | 1 | 658.06 |
| 1695 | 44% | 3 | 9 | very mild | 3 | 3 | 2 | 764.17 |
| 1700 | 92% | 1 | 35 | severe | 3 | 4 | 4 | 1264.72 |
| 1705 | 44% | 8 | 18 | mild | 2 | 3 | 4 | 692.76 |
| 1709 | 48% | 15 | 11 | very mild | 3 | 3 | 2 | 686.09 |
| 1722 | 62% | 11 | 26 | moderate | 3 | 3 | 3 | 669.07 |
| 1733 | 50% | 5 | 19 | mild | 2 | 2 | 4 | 808.89 |
| 1734 | 25% | 5 | 34 | severe | 2 | 3 | 4 | 532.59 |
| 1736 | 31% | 7 | 13 | very mild | 2 | 3 | 3 | 1040.48 |
| 1737 | 6% | 19 | 15 | very mild | 3 | 4 | 3 | 893.65 |
| 1738 | 32% | 2 | 24 | mild | 3 | 3 | 3 | 1265.39 |
| 1742 | 32% | 3 | 19 | mild | 1 | 3 | 4 | 810.22 |

#### Power-spectral density in source space

To determine the cortical origin of observed power differences between *stuttered* and *fluent* trials, we projected each participant’s epoched data to an *fsaverage* source space template (ICO 4). For each participant, we used the standard MNE-python pipeline [41] to compute a forward model based on a 1-layer boundary element model and a minimum-norm inverse model (signal-to-noise ratio = 2; loose dipole fitting = 0.2, normal orientation) using a noise covariance matrix computed from all sensors averaged over a baseline period of 300 ms across trials and determined based on a log-likelihood and cross-validation procedure [45]. This inverse model was applied to the participant’s epochs (*stuttered* and *fluent* separately, equalized in counts) using dynamic statistical parameter mapping (dSPM). Power spectral density (PSD) in source space was estimated for each trial of each condition using a multi-taper method, for times between 0.1 and 0.5 seconds after the cue and frequencies between 22 and 27 Hz (based on the TF analysis). We computed the average PSD across trials for each condition from which the normalized difference in power [(*stuttered* – *fluent)* / (*stuttered* + *fluent*)] was estimated. Each participant’s data was then morphed to a common template and entered into a cluster-based permutation test across participants (one-tailed; 1,000 permutations; N = 29) [44] with an initial *t*-threshold equivalent to *p* < 0.01 and a subsequent cluster threshold of *p <* 0.05. The location of significant clusters in cortical space was determined by projecting the anatomical parcellations defined by Glasser et al. [46] to this template and counting the number of sources within each cortical subregion (e.g., 6ma, 6mp).

#### Logistic regression analysis

We assessed the extent to which the trial-by-trial modulations of beta power in the R-preSMA, compared to baseline, predicted the corresponding trial’s outcome (stuttered, fluent). For each participant and trial, we computed the normalized difference between the beta power (22 to 27 Hz) within R-preSMA in the identified times during the cue period (0.1 to 0.5 seconds) and in an equivalent 400ms period within the baseline (−1.6 to −1.2 seconds before the cue). These trial-wise power difference values were used to predict the outcome (stuttered or fluent) of the corresponding trials using logistic regression and a cross validation procedure with 5 stratified folds [see 47 for details of this procedure]. Scores for each model were determined based on the number of correct predictions, such that if *j* outcomes are predicted correctly out of n observations (x values) the model’s score is j/n. Participants’ scores were averaged across folds and subsequently entered into a one-sample *t*-test against chance (0.5) to determine statistical significance across participants.

#### Correlation between R-preSMA beta power and percent trials stuttered

We evaluated the relationship between beta power with respect to baseline in the identified pre-SMA cluster and percent trials. We did this by fitting both linear and exponential regression models to the data and comparing the difference in explained variance (R^2^).

#### Whole-brain event-related fields analysis

In addition to the hypothesis-driven analyses in frequency space, we explored the differences in event related fields (ERFs) associated with stuttered and fluent trials using the broadband signal. MEG signals were digitally low-pass filtered at 30Hz using MNE-python’s default parameters with firwin design and epoched between −1700 ms and 1500 ms relative to the onset of presentation of the cue (white asterisk; Fig. 1). Epochs were mean-centered, linearly detrended, and equalized in counts as above so that each participant had the same number of epochs per trial type (stuttered, fluent). We applied each participant’s previously computed inverse model (see *Power-spectral density in source space* section above) to the average across trials of each condition (i.e., its ERF) and each projection was morphed to a common (fsaverage) space. Finally, we computed the normalized difference between the trial types as (stuttered-fluent)/(stuttered+fluent) before averaging across subjects. To explore differences between the trial types between the cue presentation and the go signal, we performed a series of one-sample cluster-based permutation tests [44] (1000 permutations; two-tailed) across participants (N = 29) with an initial *t*-threshold equivalent to *p* < 0.01 and a subsequent cluster threshold of *p <* 0.05 as before. Specifically, the tests were performed at the whole-brain level (i.e., using all 5124 neural sources) using a sliding window of 100 ms with a temporal overlap of 50 ms (i.e., sliding forward in increments of 50 ms). Results are reported for clusters that exceeded the *p <* 0.05 threshold.

### Data availability

All data will be made available upon the paper’s acceptance.

## Results

### Behavioral

Out of 7,964 trials, 3,166 trials were stuttered (39.75%) across participants (SD = 24.01%; MIN = 3%; MAX = 92%), and 4,798 trials (60.25%) were produced fluently. Interrater reliability for 30% of the trials, between the final author and a SLP blind to the study, yielded a Cohen’s weighted kappa of .93 (*p* < .05), indicating strong agreement [48]. Mean speech onset time was 675.84 ms (SD = 272.99). Interrater reliability for speech onset time was tested by randomly selecting 30% of the trials and calculating Cohen’s weighted kappa (*k* = 0.85, *p* < .05), indicating strong agreement [48]. Stuttered trials were associated with longer speech onset time than fluent trials (𝜷 = 0.82, t = 2.25, *p* < .05). 54.68% of independently-generated anticipated words were stuttered (SD = 37.07%; MIN = 0%; MAX = 100%), whereas 37.71% of researcher-assisted anticipated words were stuttered (SD = 24.28%; MIN = 3%; MAX = 92%). More stuttering was elicited by independently-generated than researcher-assisted words (𝜷 = 0.86, z = 6.57, *p* < .001). Due to differing abilities related to identifying anticipated words across stutterers, and also that some stutterers may not have any anticipated or feared words at this point in their lives, there was significant variability between participants in the number of independently-generated anticipated words (M = 8.51; SD = 9.26). Both word length and initial phoneme contributed to stuttered vs. fluent trials (𝜷 = −0.03, z = −2.72, *p* < .001; 𝜷 = −0.32, z = −4.07, *p* < .001, respectively) such that longer words were stuttered *less frequently*, and words starting with consonants were stuttered more frequently than words starting with vowels. The word length finding is not in line with the literature [40], but this may be because independently-generated words were generally shorter than researcher-assisted words. The initial phoneme result is in line with the literature [49,50]. Overt stuttering severity (SSI-4 score) predicted percent trials stuttered (𝜷 = 1.72, t = 3.36, *p* = .003), whereas the number of independently-generated anticipated words and stuttering impact did not (𝜷 = −0.10, t = −0.18, *p* = 0.86, 𝜷 = 0.04, t = 0.01, *p* = .99, respectively). Table 1 includes total percent trials stuttered, number of independently generated words out of 50, SSI-4 total scores and rating, responses to each subjective severity question (Q1-Q3), and speech onset time.

### Neural

#### Time-frequency in sensor space

Two participants did not attend their scheduled MEG visits (i.e., they only attended the first visit with clinical interview and diagnostic testing), thus all MEG analyses include 29 participants. A cluster-based permutation test (N = 29, 1,000 permutations; two-tailed) on the difference between stuttered and fluent trials indicated a significant difference in beta power (*p* < 0.05) with greater power in stuttered trials between 0.1 and 0.5 seconds after the cue and between 22 and 27 Hz (Fig. 2A-C). This effect remained significant (*p* < 0.05; one-tailed) after removing an outlier that was 2 SD above the mean beta power difference in the identified cluster. No significant differences in power in the theta (4 - 8 Hz), alpha (9 - 12 Hz), or low gamma (30 - 50 Hz) bands were observed. We also assessed the relationship between beta power differences within the reported cluster (22 to 27 Hz and 0.1 to 0.5 seconds) and stuttering severity (indexed by SSI-4 scores) and found a significant positive correlation (Spearman rho = 0.33, *p* < 0.05; Fig. 2D).

**Figure 2.**
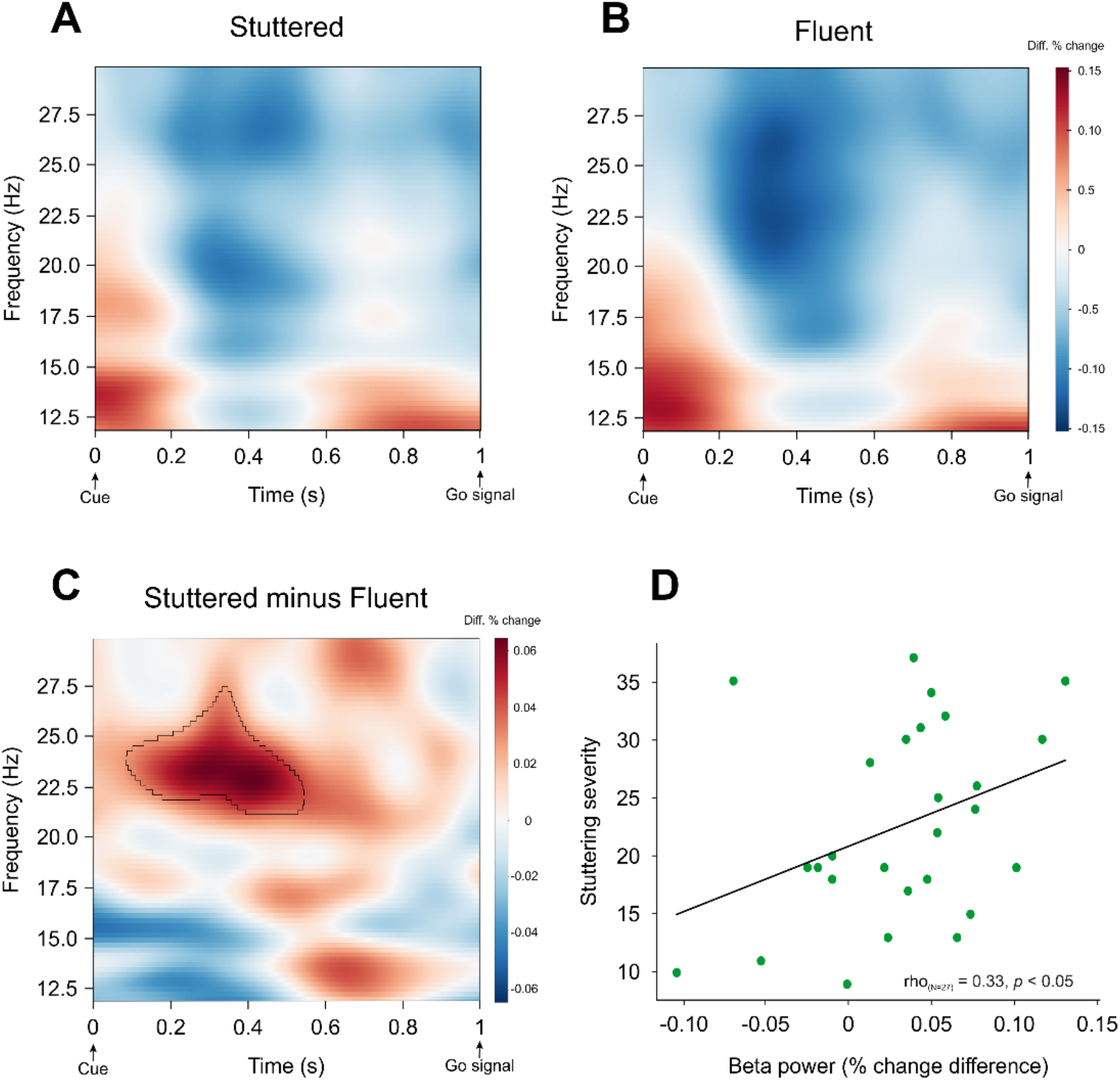
Time frequency analysis. A-C. Greater beta power prior to stuttered compared to fluent speech. Time-frequency plots for stuttered (**A**), fluent (**B**), and the difference (stuttered minus fluent; **C**). The significant beta band cluster in panel C is highlighted (black contour). Significance was determined via a cluster-based permutation test (*p* < 0.05; one-tailed; 1,000 permutations) across participants (N = 29). The time-frequency analysis window spanned 1 second between the cue (white asterisk; Fig. 1) and the go signal (green asterisk). Zero (0) is the time when the cue was presented; Time 1 is when the go-signal was presented. **D. Correlation between beta power and stuttering severity.** Scatterplot of the relationship between SSI-4 scores (a proxy for stuttering severity) and beta power (22 to 27 Hz, 0.1 to 0.5 seconds after the cue) with Spearman rho values of the correlation.

There was also a significant positive correlation between beta power differences and participants’ number of independently-generated words (Spearman rho = 0.34, *p* < 0.05).

#### Power-spectral density in source space

To determine the cortical origin of beta power differences between stuttered and fluent trials, we computed the normalized difference in power [(*stuttered* – *fluent)* / (*stuttered* + *fluent*)] from each participant’s average power spectral density for each condition in source space. A cluster-based permutation test (N = 29; 1,000 permutations; one-tailed) across participants using participants’ source projected data (the entire cortical surface morphed to a common *fsaverage* space) spanning 0.1 to 0.5 seconds after the cue and a 22 - 27 Hz frequency range yielded significant differences (*p* < 0.05) in a single cluster comprising the right anterior and medial SMA (i.e., preSMA; Fig. 3). The precise location of the cluster in cortical space was determined by quantifying the number of sources in the cluster falling within each label of the Glasser et al.’s [46] cortical atlas. The largest number of sources corresponded to label 6ma (medial anterior) in the right hemisphere (i.e., to the R-preSMA). No sources corresponded to the right SMA proper (label 6mp [medial posterior]).

**Figure 3.**
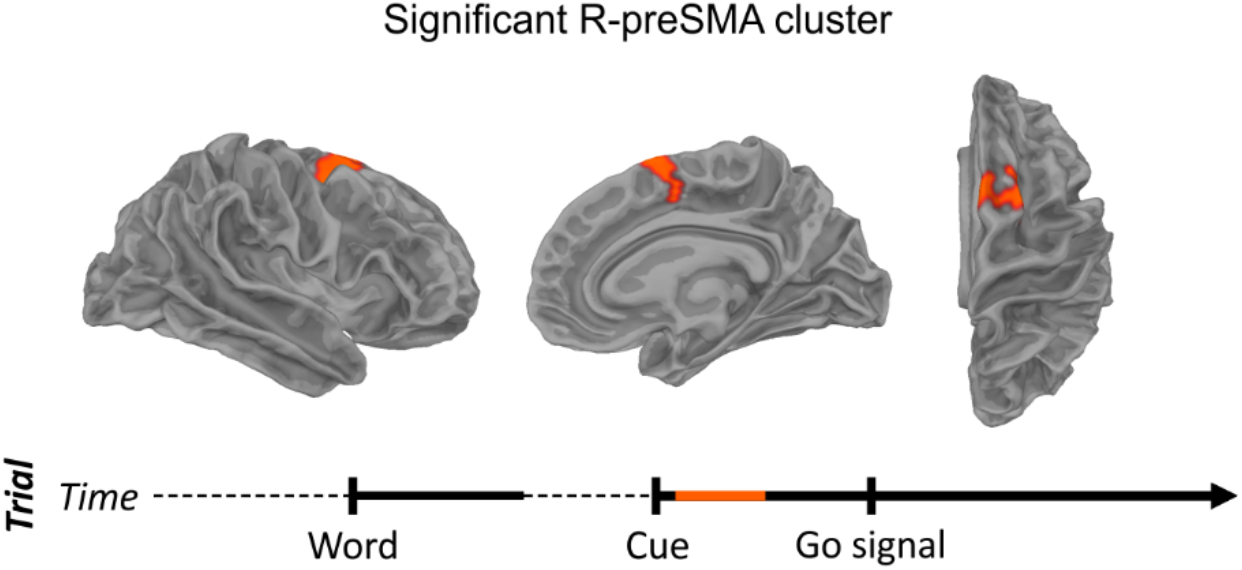
Greater beta power in the R-preSMA prior to stuttered speech. Significant cluster shown in a lateral (**left**), medial (**middle**), and axial (**right**) views. Significance was determined via a cluster-based permutation test across participants using each participant’s power spectral density estimates in source space (5124 sources) averaged over frequencies 22 – 27 Hz and times 0.1 to 0.5 sec after the cue, with a *t*-threshold of *p* < 0.01 and a subsequent cluster threshold of *p <* 0.05. Timeline shows pertinent time window of interest in orange.

Importantly, although motor beta suppression was present, reflecting typical speech planning or replanning processes, no difference was found between stuttered and fluent trials (Fig. S1).

#### Relationship between R-preSMA power and stuttering outcome

Based on the above results, we next evaluated the extent to which each subject’s trial-by-trial beta power modulations in the R-preSMA, with respect to baseline, predicted the trial’s outcome (fluent, stuttered). For each participant, we used a 5-fold cross-validation procedure and logistic regression to predict each trial’s outcome based on that trial’s beta power compared to baseline. A one-sample *t*-test across subjects against chance (0.5) on each subject’s logistic regression model scores (averaged across folds) indicated that the trial-by-trial beta power in the R-preSMA predicted whether subsequent speech would be stuttered (*t*(28) = 3.56, *p* < 0.001).

We next assessed the relationship between beta power in the R-preSMA cluster and percentage of stuttered trials. We found a positive correlation between each participant’s mean beta power across trials (relative to baseline) and percent trials stuttered (Spearman rho = 0.5, *p* < 0.01; Fig. 4), indicating that increases in beta power in the R-preSMA are associated with more stuttering. The superior fit of an exponential model (R^2^ = 0.57; Fig. 4, right) over a linear regression model (R^2^ = 0.22; Fig. 4, left) indicates that the percentage of trials stuttered increased exponentially with beta power increases in the R-preSMA.

**Figure 4.**
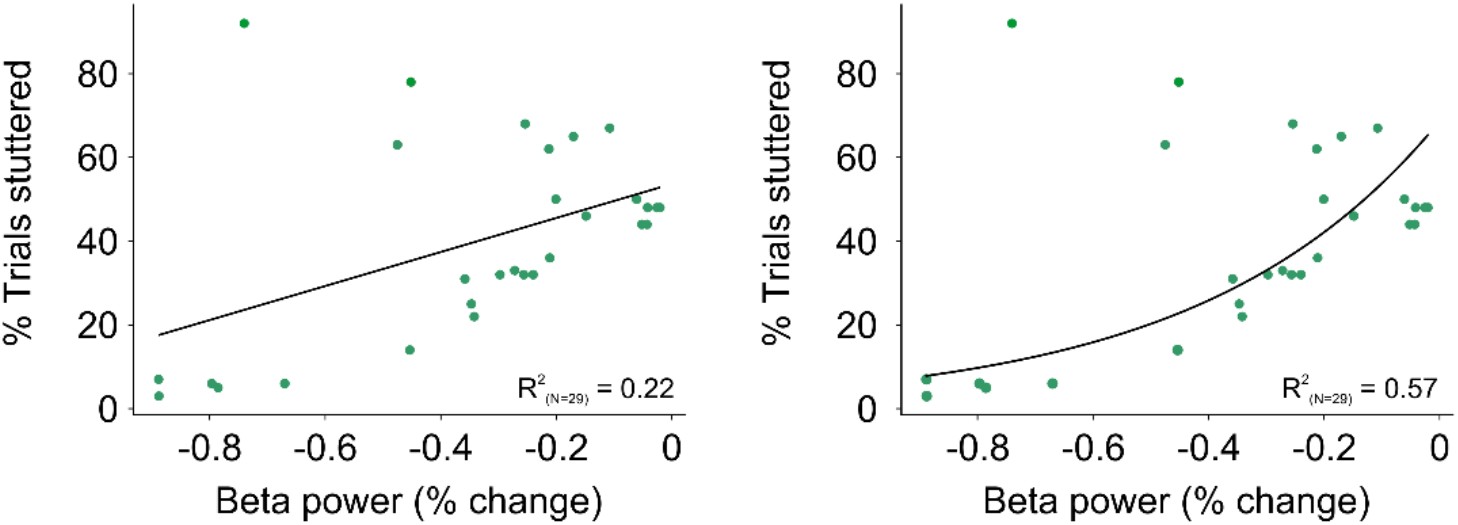
Relationship between pre-speech R-preSMA beta power and percentage of trials stuttered. Left: The plot shows a positive linear relationship between beta power changes compared to baseline in the R-preSMA and percentage of trials stuttered. **Right:** Exponential fit to the same data as the left plot. The greater variance explained (R^2^) for the exponential fit suggests that the percentage of trials stuttered grows exponentially as beta power increases.

In addition to the hypothesis-driven analyses discussed in this section, we conducted a whole-brain event-related fields analysis of the differences between stuttered and fluent trials (Fig. 5). A similar R-preSMA cluster emerged between 200 ms and 300 ms after the cue in line with the time-frequency and power spectral density results (Fig. 5 inset). R-preSMA activity was followed by significant differences in the R-DLPFC 500 ms after the cue as well as in the left SMA and posterior superior temporal gyrus.

**Figure 5.**
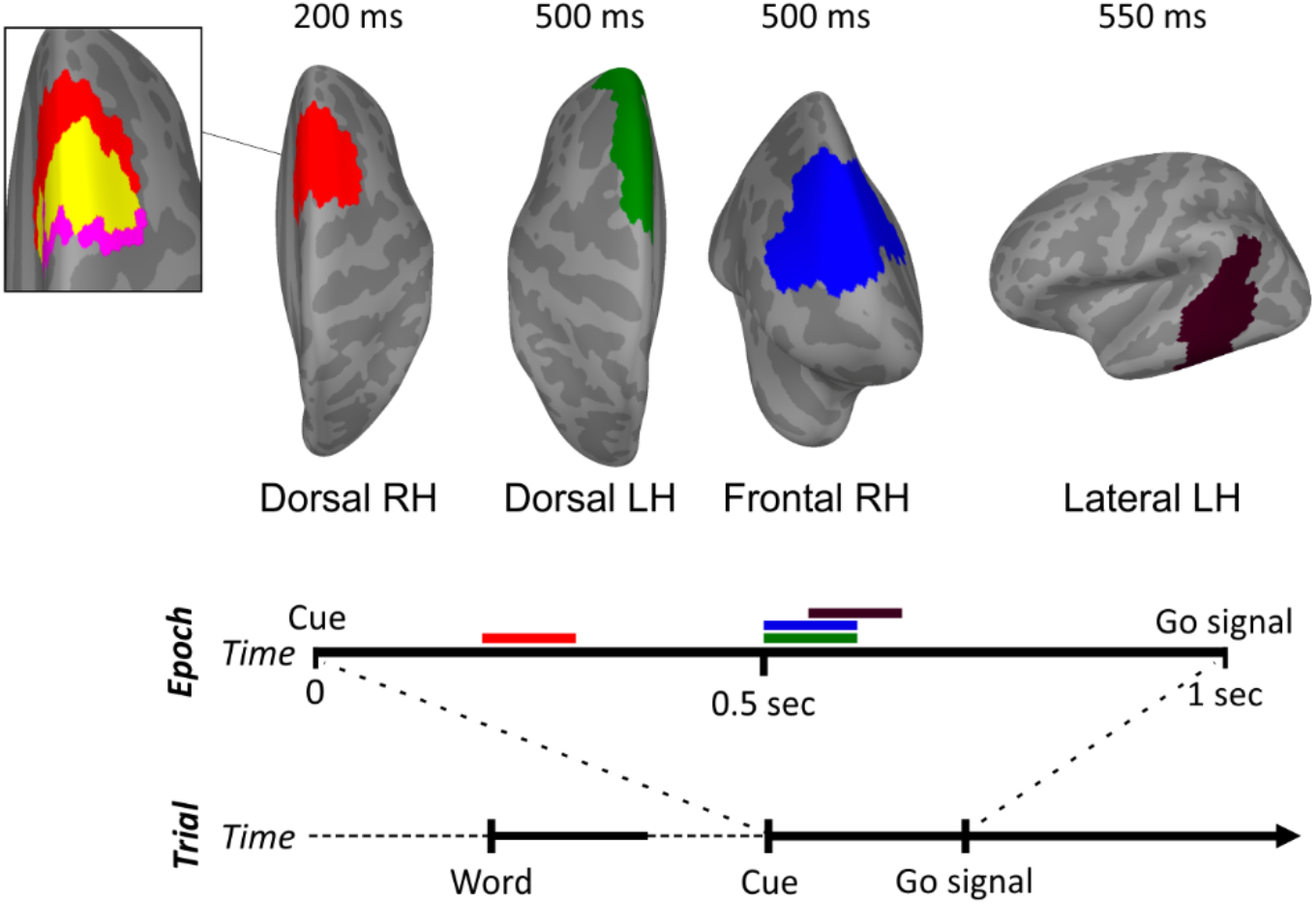
Whole-brain ERF analysis results. The analysis yielded four distinct clusters with differential neural activity for stuttered compared to fluent trials (*p* < 0.05). Between 200 and 300 ms after the cue, there was divergent activity for stuttered vs fluent trials in the right dorsal frontal cortex (red cluster), in a cluster largely overlapping in time and spatial location that identified in the PSD analysis (inset: R-preSMA cluster from Fig. 3 [magenta]; overlap [yellow]). This was followed by divergent activity for stuttered trials between 500 and 600 ms in the left superior frontal gyrus (green cluster) and in the R-DLPFC (blue cluster). There was also divergent activity in the left posterior superior and middle temporal cortex (dark violet cluster) between 550 and 650 ms. The timeline of events in the experimental trial and the temporal segment corresponding to the epoched data (0 to 1 second after cue presentation) are shown below the clusters. The times in which significant differences were found are indicated by horizontal bars with the color of each cluster. RH: Right Hemisphere. LH: Left Hemisphere.

## Discussion

This study aimed to determine neural markers preceding stuttered speech. We overcame the significant challenge of eliciting stuttering during neuroimaging by implementing a relatively novel procedure [36,37] resulting in the largest and most balanced neural investigation of stuttered vs. fluent speech to date. The premise of the study was that stutterers are generally averse to producing stuttered speech and therefore will exhibit an inhibitory control response prior to producing overtly stuttered versus fluent words. Our previous work using fNIRS (measuring a slow hemodynamic response) pointed to a proactive inhibitory control response in the seconds leading up to the production of stuttered vs. fluent (and anticipated vs. unanticipated) words. Here, we leveraged the high temporal resolution of MEG to test the hypothesis that a reactive inhibitory control response is triggered when a cue signals the impending requirement to produce a word likely to be stuttered.

The primary finding of this study is that, compared to fluent speech, stuttered speech is preceded by increased beta power in the R-preSMA following a cue indicating that speech is imminent. Given its cortical origin and timing relative to the cue, this neural response is consistent with reactive inhibitory control or a freezing-like response [33]. Indeed, reactive inhibitory control is associated with increased beta power in the R-preSMA in response to stop signals [26,32,51]. During classic stop signal tasks, a go signal is displayed in each trial, prompting participants to perform a specific action (e.g., pressing a button). For a small proportion of trials, a stop signal is introduced shortly after the go signal, requiring the individual to stop their automatized response. Elevated activation in the R-preSMA is understood as a reaction to the stop signal because of its automatic and fast nature (typically occurring within 120ms after the stop signal) [31,52,53]. A similar neural response is elicited in go/no-go tasks in which a habitual response to go signals needs to be inhibited in a small proportion of (no-go) trials [26]. In the current study we reasoned that a cue signaling the requirement to imminently produce a word likely to be stuttered would act as an implicit stop or no-go signal, and thus result in a similar reactive and automatic inhibitory control response because the stutterer does not want to stutter overtly. Our data are consistent with this hypothesis.

While we observed greater activity in the R-preSMA, we did not find elevated activity in the R-IFG. Elevated activity in the R-IFG has been associated with reactive inhibitory control [54–56] and is also a common finding in studies comparing the fluent speech of stutterers and non-stutterers [e.g., 30,57,58]. It is unclear why the current result was restricted to the R-preSMA, however, it may have to do with the stuttering state. While the R-IFG is considered to be a neural signature of stuttering [59,60], this is primarily the case when stutterers are speaking fluently. In addition, previous studies assessed activation during or time-locked to speech execution compared to pre-speech and time-locked to a cue as in the current study. It is likely, for example, that both the aversion to stuttered speech (prior to execution) and consequent motor inhibition are absent prior to fluent speech. It may be that previous studies showing elevated activation in the R-IFG point to compensatory processing which facilitates fluent speech in stutterers which is not present when speech is stuttered. In addition, the preSMA in conjunction with the STN [61–63] (and not the R-IFG) has been implicated in other movement-related disorders. For example, reduced preSMA-STN connectivity is associated with freezing of gate in Parkinson’s disease when these individuals walk through a doorway which was also associated with longer footstep latency. Given that the relationship between the R-IFG and preSMA in action-stopping remains unclear, future studies can probe this relationship for example by focusing on disorders like stuttering to help determine the respective roles of the R-preSMA and R-IFG.

Reactive inhibitory control differs from proactive inhibitory control which is forward-looking and more deliberate—for example, the speaker has more time to reflect on the sense that upcoming speech will be stuttered. Neurally, these types of inhibition are thought to be mediated through distinct pathways (hyperdirect and indirect BGTC pathways, respectively) and, consequently, at different rates (faster and slower, respectively). Proactive inhibitory control could occur seconds or longer before an expected action, whereas reactive inhibitory control would immediately follow some kind of internal or external cue associated with halting the currently planned action. Still, it is likely that proactive and reactive inhibitory control are related and overlapping processes [26,64]. In the context of stuttering, Jackson et al. [17] found evidence for proactive inhibitory control preceding the production of words likely to be stuttered. In that study, there was a five-second delay between the time when participants saw the word that they would have to produce and the go signal [17]. Although the current experiment was designed to reveal a reactive inhibitory response, it is possible that proactive control was similarly initiated soon after the words (which were subsequently stuttered) were presented, and sustained until the word was produced. This is in line with the activity found in the R-DLPFC in the current study, albeit occurring after the inhibitory control response to the cue. Alternatively, R-DLPFC activity could reflect error monitoring responses to a recognition that something has gone awry during the speech planning process [17]. Future work can further probe the nature of cognitive control involvement preceding overtly stuttered speech, and also potential differences between proactive and reactive inhibitory control in stutterers by using tasks specifically designed to tease apart proactive and reactive inhibitory control, if that were possible.

Generally, our findings are consistent with proposals that hyperactive inhibitory control in the right hemisphere BGTC loop is involved in stuttering [29–31,65]. These largely speech motor-based accounts propose that inhibitory control could cause the interruptions in speech that characterize overt stuttering (i.e., blocks, prolongations, repetitions) by suppressing the initiation or sequencing of speech motor movements [29]. This could occur by interfering with the forward modeling of sensory predictions required for online speech motor control (in line with [12]). Deficient forward modeling would contribute to inaccurate auditory feedback, consistent with Chang and Guenther’s account [66]. Our results offer some support for these ideas, showing divergent activity preceding stuttered vs. fluent productions in the left posterior temporal cortex and SMA, i.e., regions associated with the processing of auditory feedback [67]. However, proponents of the hyperactive inhibitory control account do not specify why reactive inhibitory control would arise in the first place, suggesting that it may occur due to random neural fluctuations in the right hemisphere BGTC loop. While this is plausible, it is difficult to find a reason for why this would be the case. Our account is that during development, stutterers learn to anticipate stuttering events by associating certain words or sounds with instances of stuttering and their negative consequences [21,68]. We argue that when stuttering on a particular word or sound is anticipated, its production is inhibited because the speaker does not want to stutter to avoid these negative consequences. This is consistent with our findings of delayed speech onset time for stuttered words, and that independently-generated anticipated words are related to both more stuttering and greater beta power than researcher-assisted words because independently-generated words are more likely to trigger these negative associations. Once these associations are in place, inhibitory control may act proactively, for example, when the speaker knows they will have to produce a word likely to be stuttered (as in [17]), or reactively, triggered by a cue (implicit or explicit) signaling the requirement to imminently produce the anticipated word. Importantly, we believe that stuttering can occur in the absence of elevated inhibitory control, particularly during spontaneous speech when there is either not enough time to retrieve these associations (i.e., to anticipate stuttering) or the speaker’s attention is not directed towards the speech production process. Critically, the paradigm in the current study created a setting in which it was likely that if a word was stuttered, it was also anticipated. As a result, elevated inhibitory control was also likely.

While the current results are consistent with a reactive inhibitory control response in the R-preSMA, it is necessary to consider alternative interpretations. For example, the suppression of beta oscillations is commonly reported in electrophysiological studies of motor execution [e.g., 69,70]). Although we attempted to minimize co-occurrence of motor beta suppression and reactive inhibition in our task by separating word presentation and cue by a jittered interval, our data indicate that some planning or re-planning took place during the window of interest after the cue (Fig. S1). It could thus be that stuttered trials exhibit less beta suppression than fluent trials. However, motor beta suppression in our task localized, as expected [e.g., 69,71,72], to bilateral speech motor cortex and did not distinguish between stuttered and fluent trials, in line with Mersov et al. (2018). The transient nature of the response and its timing relative to the cue suggest that it is also unlikely that the difference in beta for stuttered vs fluent trials we report reflects the MEG counterparts of lateralized readiness potential or contingent negative variation responses, which are typically slow ramping and more closely aligned to motor execution [73,74].

The current findings, as well as other recent work [17], have important implications for neuromodulation as a possible complement to existing behavioral therapy. Transcranial direct current stimulation (tDCS) is starting to be applied in stuttering research [75–77], albeit with modest results. For example, Garnett et al. [76] tested the impact of anodal tDCS on overt severity in 14 adult stutterers, and while they did not find significant effects on overt stuttering severity, they found that the atypically strong association between overt severity and right thalamocortical activity was attenuated after tDCS, especially in severe stutterers. It may be that the modest effects reported to date are due to an exclusive focus on the speech network. Future neuromodulation studies can target, for example, proactive (R-DLPFC) and reactive inhibition (R-preSMA) to test whether forward-moving speech is facilitated by reducing the putative interference from hyperactive inhibitory control on speech production.

### Limitations

There were two primary limitations of this study. First, spatial precision of MEG is limited and therefore we cannot be certain that activation emanated from the R-preSMA specifically. However, we argue that the theoretical foundation for a right pre-SMA response is strong, not only in light of the action-stopping literature [26,31,52] and our task design but also of our previous work on anticipation [17,18,27] and the experience of people who stutter [3,4]. Second, we did not directly measure anticipation during the MEG experiment primarily because there is currently no feasible way to do so. This is one of the motivations of the current study, to better characterize the neural processes leading up to stuttered vs. fluent speech. We decided not to ask participants during the experiment whether anticipation occurred because this could 1) change the outcome of whether the word will actually be stuttered—if participants are asked at the beginning of the trial; or 2) be influenced by whether the word was stuttered— if participants are asked at the end of the trial. Nonetheless, it is critical to highlight that adult stutterers predict upcoming stuttering with high accuracy (>90%) [22–24] particularly when there is a delay between stimulus presentation (i.e., when the stutterer is made aware of the word to be said) and speech execution [17]. Therefore, we can be confident that participants anticipated prior to stuttered trials but not prior to fluent trials.

## Conclusion

This study is the largest and most balanced neurofunctional investigation of stuttered vs. fluent speech to date. We found elevated beta activity in the R-preSMA preceding stuttered vs. fluent speech which was related to stuttering severity and proportion of trials stuttered. Stuttered trials were also associated with delayed speech onset. These neural and behavioral results suggest a reactive inhibitory control response to cues that signal to stutterers that they will soon have to produce a word that will likely be stuttered, which is consistent with previous accounts of freezing in stuttering. A post-hoc analysis also pointed to a relationship between stuttering anticipation and this reactive inhibitory control response such that words that were associated with greater anticipation (i.e., independently-generated vs. researcher-assisted anticipated words) were related to greater beta power and more stuttering. This exploratory finding suggests that anticipation precedes reactive inhibitory control such that the inhibitory response is triggered because the speaker can predict and is averse to upcoming overt stuttering. While elevated inhibitory control is likely not the cause of stuttering events, it may contribute to stuttering events by interfering with speech motor control (e.g., thinking consciously about an automatic process), and ultimately, make it more difficult for stutterers to move forward during speech. Broadly, the current results support a neurocognitive perspective of stuttering events by characterizing cognitive processes that precede overtly stuttered speech. Future work should continue to investigate the role of inhibitory control in stuttering events as well as explore the possibility of targeting the neural regions underlying hyperactive inhibitory control to facilitate speech production in stutterers.

## Author contributions

Joan Orpella: Conceptualization: Lead; Formal Analysis: Lead; Investigation: Lead; Methodology: Lead; Writing: Lead. Graham Flick: Conceptualization: Equal; Formal Analysis: Supporting; Investigation: Equal; Methodology: Supporting; Writing: Supporting. M. Florencia Assaneo: Conceptualization: Equal; Formal Analysis: Supporting; Methodology: Supporting; Writing: Supporting. Liina Pylkkänen: Conceptualization: Supporting; Writing: Supporting. David Poeppel: Conceptualization: Supporting; Writing: Supporting. Eric S. Jackson: Conceptualization: Lead; Formal Analysis: Lead; Funding Acquisition: Lead; Investigation: Lead; Methodology: Lead; Project Administration: Lead; Writing: Lead.

## Supplementary Material

**Table S1.** Number of trials per participant and condition (Stuttered, Fluent) after equalizing the number of trials (see Methods).

| ID | Num. of trials<br>per condition |
| --- | --- |
| 1607 | 115 |
| 1609 | 52 |
| 1610 | 100 |
| 1611 | 56 |
| 1613 | 51 |
| 1614 | 85 |
| 1615 | 104 |
| 1621 | 83 |
| 1634 | 41 |
| 1636 | 102 |
| 1637 | 8 |
| 1675 | 145 |
| 1678 | 7 |
| 1679 | 13 |
| 1681 | 31 |
| 1686 | 100 |
| 1687 | 95 |
| 1690 | 83 |
| 1695 | 124 |
| 1700 | 19 |
| 1705 | 75 |
| 1709 | 137 |
| 1722 | 109 |
| 1733 | 143 |
| 1734 | 72 |
| 1736 | 80 |
| 1737 | 16 |
| 1738 | 93 |
| 1742 | 90 |

**Figure S1.**
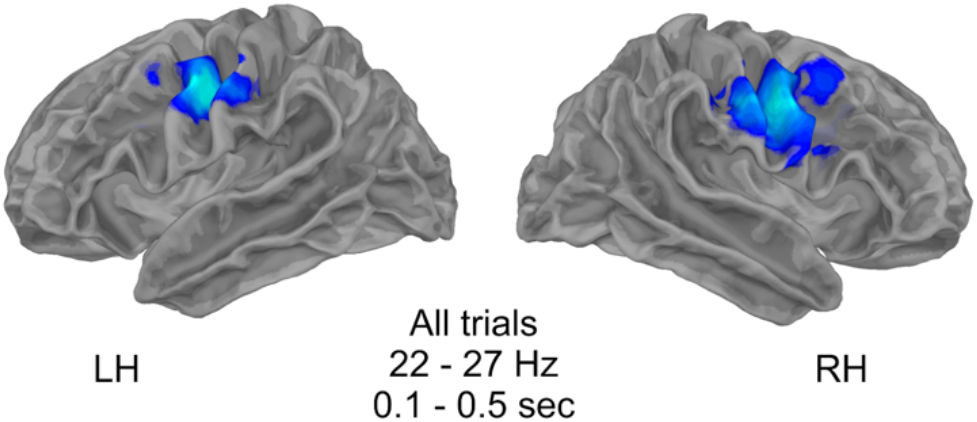
Motor beta suppression over speech motor cortex preceding speech production. No significant differences were found between stuttered and fluent trials.

